# Parkinsonism Reversal and Dopaminergic Resilience: Lessons from a Rotenone-induced Parkinson’s Disease Model

**DOI:** 10.64898/2025.12.20.695709

**Authors:** Prince Joshi, Fan Fan, Xuelin Lou

## Abstract

Parkinson’s disease (PD) is a progressive neurodegenerative disorder characterized by profound loss of dopaminergic (DA) neurons, yet the underlying mechanism remains incompletely defined. Mitochondrial toxins can induce acute degeneration of DA neurons and Parkinsonism-like phenotypes in animal models, and epidemiological studies have linked pesticide exposure to increased PD risk; however, the long-term effects of pesticide exposure remain elusive. Here, we examined both the acute and long-term effects of rotenone exposure in mice to understand PD onset, progression, and recovery. A 21-day regimen of rotenone intraperitoneal injections (2.5 mg/kg/day) induced robust Parkinsonism-like deficits by the 4^th^ week, including impaired locomotion, increased anxiety-like behaviors, and deficits in motor balancing and coordination. These behavioral abnormalities were accompanied by pronounced reduction in tyrosine hydroxylase (TH) expression and selective loss of DA neurons in the substantia nigra pars compacta (SNc). Unexpectedly, these functional impairments fully resolved by 12 months, and rotenone-treated mice behaved equally well as age-matched controls. In parallel, the TH expression and DA neuron density in SNc were restored to control levels. Together, these longitudinal results demonstrate that chronic rotenone injection induces robust but reversible Parkinsonism in the acute phase, with limited long-term consequence on Parkinsonism upon toxin cessation. These findings contrast with the prevailing view that environmental pesticide exposure irreversibly drives PD and instead they reveal a substantial resilience and adaptive capacity of the nigrostriatal dopaminergic system in vivo.

**Highlights:**

- Chronic rotenone exposure in mice induces robust Parkinsonism-like behaviors in both motor and non-motor domains.
- SNc dopaminergic neurons are reduced by ∼40% at the acute phase within 4 weeks.
- Both motor and non-motor deficits are fully recovered by 12 months.
- PD-like pathological changes in the SNc are resolved by 12 months.

**Infographics:** 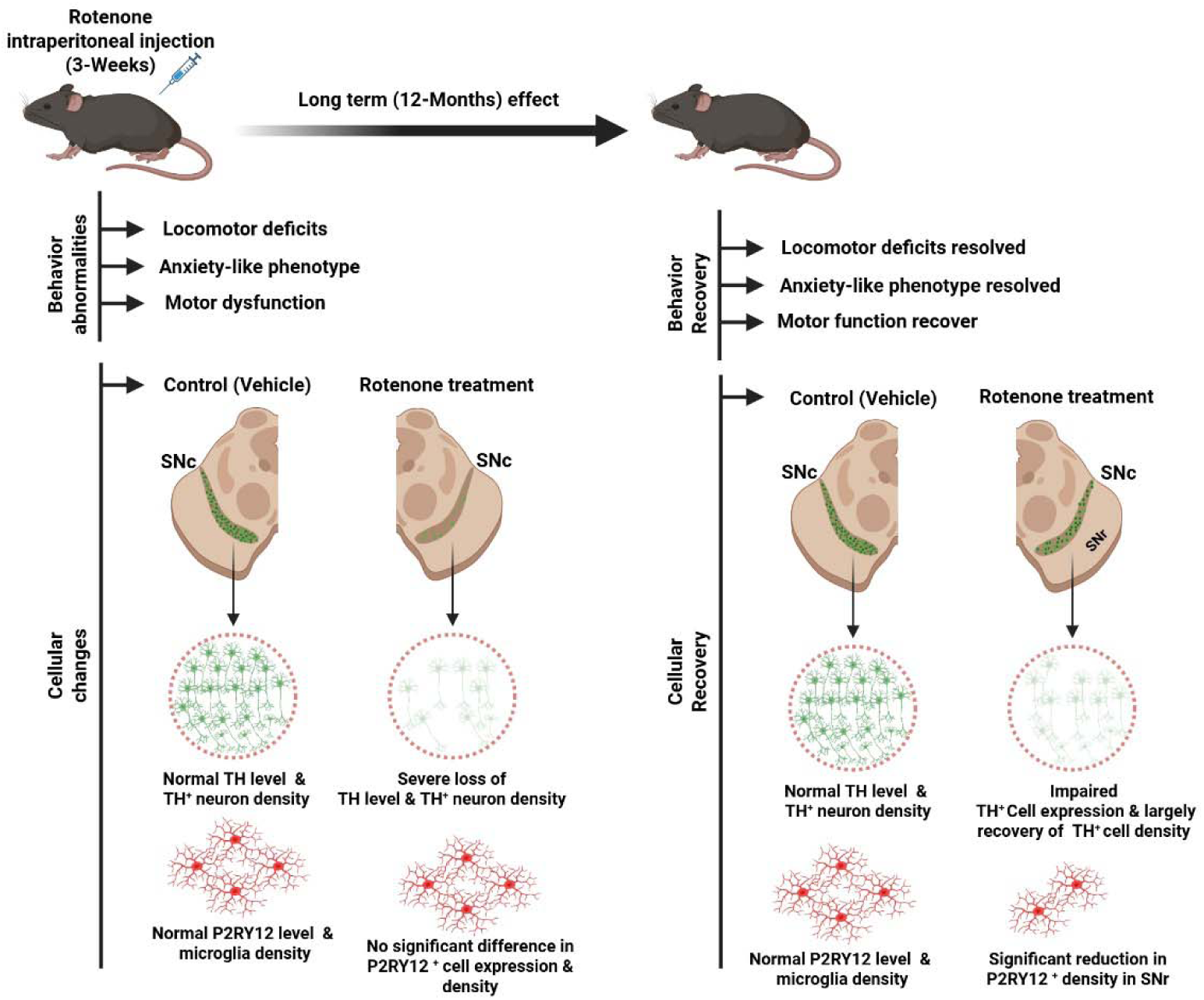

## Introduction

Parkinson’s disease (PD) is the second most common neurodegenerative disease, but current treatments are limited to symptomatic relief only, and none of them halt or slow disease progression. Clinically, PD exhibits cardinal motor symptoms, including akinesia, bradykinesia, tremor, rigidity, and postural instability, as well as prominent non-motor symptoms such as anxiety, constipation, hyposmia, and sleep disturbance^1,2^. These manifestations arise primarily from the progressive loss of dopaminergic (DA) neurons in the substantia nigra pars compacta (SNc) and consequent depletion of dopamine in the caudate putamen. Despite decades of research, PD pathogenesis remains incompletely understood; it may involve a complex interplay between genetic susceptibility and environmental factors converging on key processes such as misfolded α-synuclein, overwhelming metabolism, and chronic inflammation^2^. Across both sporadic and familial forms of PD, mitochondrial dysfunction has emerged as a central player. Impaired mitochondrial complex I, elevated oxidative stress, and bioenergetic failure are consistently observed in animal models and postmortem PD brains. In line with this, exposure to pesticides containing mitochondrial toxins—most notably rotenone and paraquat—has been associated with an increased risk of PD^3–5^, but a definitive causal relationship remains uncertain^3,4^. Addressing causality requires longitudinal studies that are not feasible in human populations, underscoring the importance of well-controlled animal models.

Rotenone, a widely used pesticide and potent mitochondrial complex-I inhibitor, provided early experimental support for the mitochondrial hypothesis of PD. Chronic rotenone infusion in rats via jugular vein cannula for 1–5 weeks can induce selective SNc degeneration, synuclein aggregation, and hypokinesia^6^. Since this seminal work, rotenone has been increasingly used to model PD-like pathology and to evaluate candidate PD drugs^7–9^. Compared with other commonly used toxins, such as 6-hydroxydopamine (6-OHDA), N-methyl-4-phenyl-1,2,3,6-tetrahydropyridine (MPTP), and its metabolite (1-methyl-4-phenylpyridinium (MPP^+^), rotenone models offer distinct advantages. The latter toxins rely on dopamine transporter-dependent uptake to induce selective dopaminergic lesions, effectively modeling late-stage PD within hours or days of administration^10–12^ and thus limiting their ability to capture progressive neurodegeneration or α-synuclein aggregation. In contrast, rotenone exposure better mimics the insidious process onset of PD and reproduces more disease-relevant processes, including mitochondrial impairment^13^, nigrostriatal dopaminergic loss^6,14,15^, α-synuclein inclusions akin to Lewy bodies^6,14,16^, and neuroinflammation^17–19^. Notably, transient rotenone exposure in rats has been reported to induce progressive posture instability and delayed α-synuclein aggregation after 9 months of exposure^16^.

While rotenone rat models have been well established for decades^6,7^, mouse rotenone models have gained attention only more recently, with different routes of exposure reported ^9,20,21^. These models exhibit substantial variability depending on experimental conditions, including exposure route, age, and sex, with males producing more robust phenotypes than females^9,22,23^. Compared with rat models, outcomes in mice are less consistent; even identical oral exposure paradigms have yielded conflicting results^14,24^. Although these rotenone rodent models are valuable in vivo systems for interrogating PD pathogenesis and therapeutic strategies, most studies focused on short-term outcomes, leaving the long-term consequences of rotenone exposure unexplored.

To address this knowledge gap, we systematically evaluated the effectiveness and long-term reversibility of PD-like behaviors and associated neuropathology induced by a chronic systemic rotenone exposure—three weeks of intraperitoneal injection (IP). We show that rotenone exposure induces a robust Parkinsonism during the acute phase shortly after exposure cessation. Unexpectedly, however, both behavioral deficits and dopaminergic pathology recover to control levels by 12 months. These findings reveal a previously underappreciated capacity for adaptive compensation and resilience within the dopaminergic system in vivo, with potential implications for better understanding PD progression and recovery.

## Results

### Chronic systemic rotenone exposure in mice impairs locomotion at the acute phase

The age-matched male mice were treated with rotenone and vehicles in parallel for 21 days, with their behavior changes monitored. Before experiments, rotenone was dissolved in 100% DMSO first and then diluted in medium-chain triglyceride Miglyol 812, injected intraperitoneally (2.5 mg/kg bodyweight per day). Mice receiving rotenone were referred to as the PD model; mice receiving only vehicle in parallel were used as controls. Open field tests were performed before the treatments and within the 4^th^ week of treatments, referred to as the acute phase. The results demonstrated a strong impairment in locomotion in rotenone-treated mice (Figure 1). These mice exhibited significant reduction of total travel distance, movement speed, and mobile time, and the increased immobile time and freezing time (*n* = 12 mice for vehicle group; *n* = 15 mice for PD model group, two-tailed Welch’s *t*-test). Thus, systemic rotenone IP exposure impairs mouse locomotor function at the acute phase, mimicking motor deficits in human PD.

**Figure 1.**
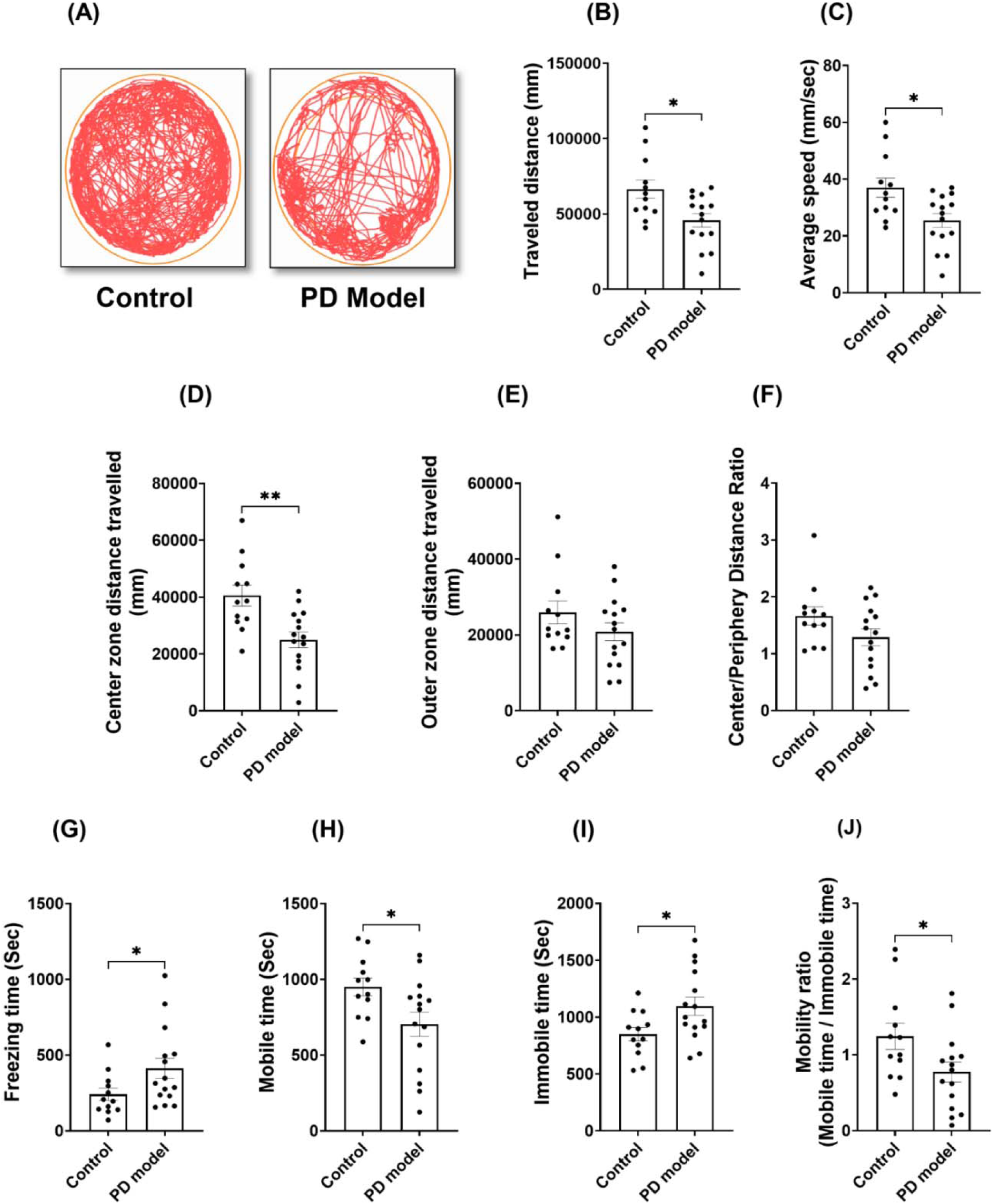
Intraperitoneal (IP) rotenone exposure induces locomotor deficits at the acute phase. (A) Representative movement tracks in open field tests from control and PD model (rotenone-treated) mice, which were measured after 3 weeks of treatment. (B-D) Reduction in the total distance traveled (B), average speed (C), and center-zone distance traveled (D) was observed in rotenone-treated mice. (E-F) Intact outer zone distance traveled (E) and center to periphery distance ratio (F). (G) Freezing time. (H-I) Mobile and Immobile time. (J) Mobility ratio. Data were shown as mean ± SEM. Individual points represent single animals. Statistical analyses were performed using two-tailed Welch’s *t*-tests. Sample sizes: (control *n* = 12, PD model *n* = 15). Statistical significance is indicated as *p* < 0.05 (*) and *p < 0.01 (*\*\*).

### Rotenone systemic exposure induces anxiety

In the open field tests above, we observed a significant reduction of travel distance selectively in the center (but not outer) zones (Figure 1D-E), a typical sign of anxiety-like behaviors. To verify this abnormality, we performed the elevated plus maze (EPM) tests (Figure 2), another assay for evaluating anxiety behaviors, at the 4th week after rotenone exposure. Rotenone-treated mice showed a significant reduction of total travel distance (Figure 2B), consistent with the impaired locomotion in the open field results (Figure 1D). Importantly, these mice displayed a significantly lower number of open-arm entries, and less time and distance traveled in the open-arm (Figure 2C-E). These results suggest that rotenone induced non-motor deficit at this acute stage, a psychiatric symptom often observed in PD patients before motor dysfunction.

**Figure 2.**
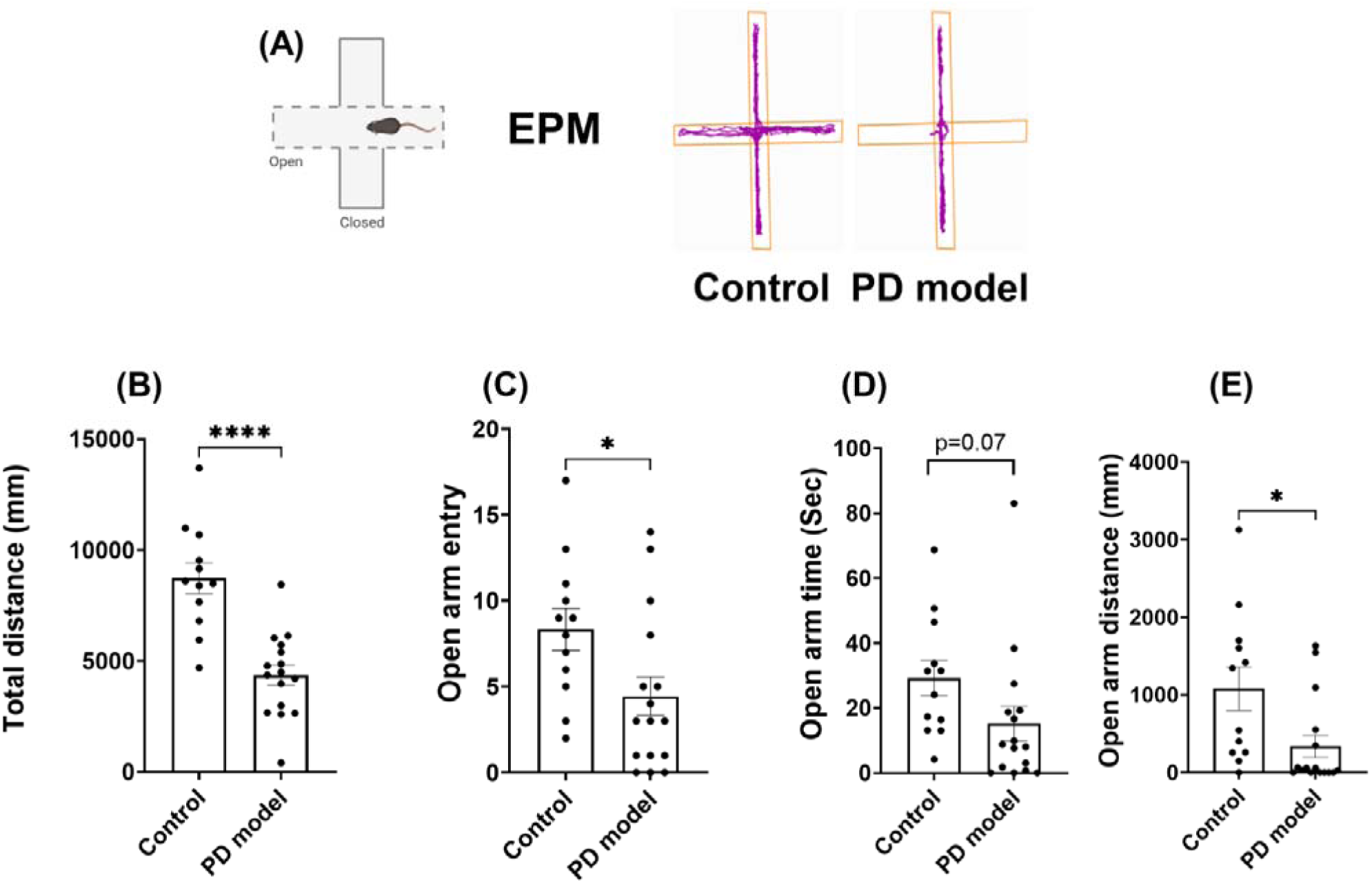
Intraperitoneal rotenone exposure increases anxiety-like behaviors at the acute phase. (A) Representative tracking plots of mouse trajectories in the elevated plus maze (EPM). (B) total distance traveled in the EPM. (C) open-arm entry. (D) open-arm time. (E) open-arm distance. Experiments were performed 3 weeks after treatments. Data are shown as mean ± SEM. Individual points represent single animals. Statistical analyses were performed using two-tailed Welch’s *t*-tests. Sample sizes: Control *n* = 12, PD model *n* = 16. Statistical significance is indicated as *p < 0.05 (*), and p < 0.0001 (****).* IP, intraperitoneal.

### Rotenone exposure disrupts motor balance, coordination, and motor learning

Next, we examined our PD model for the changes in motor balance and coordination. Male mice were trained and tested using the rotarod task right after the day of completing chronic rotenone exposure (Figure 3). The results showed rotenone-treated mice performed significantly worse than controls, with a significantly shorter latency to fall, suggesting severely impaired motor balance and coordination during the task (Figure 3A). Further, while the control group improved their performance significantly 3 weeks later as compared to the initial level (baseline), the rotenone-treated group showed worse performance (Figure 3B-C). These distinct time-dependent changes between the two groups may stem from different motor learning, which was impaired in the rotenone-treated group. Additional Y-maze tests showed no difference between the two groups (Figure 3D), indicating intact spatial memory in the rotenone group. Together, these data suggest that rotenone treatment significantly impairs motor balance and coordination, as well as preferentially disrupts motor learning.

**Figure 3.**
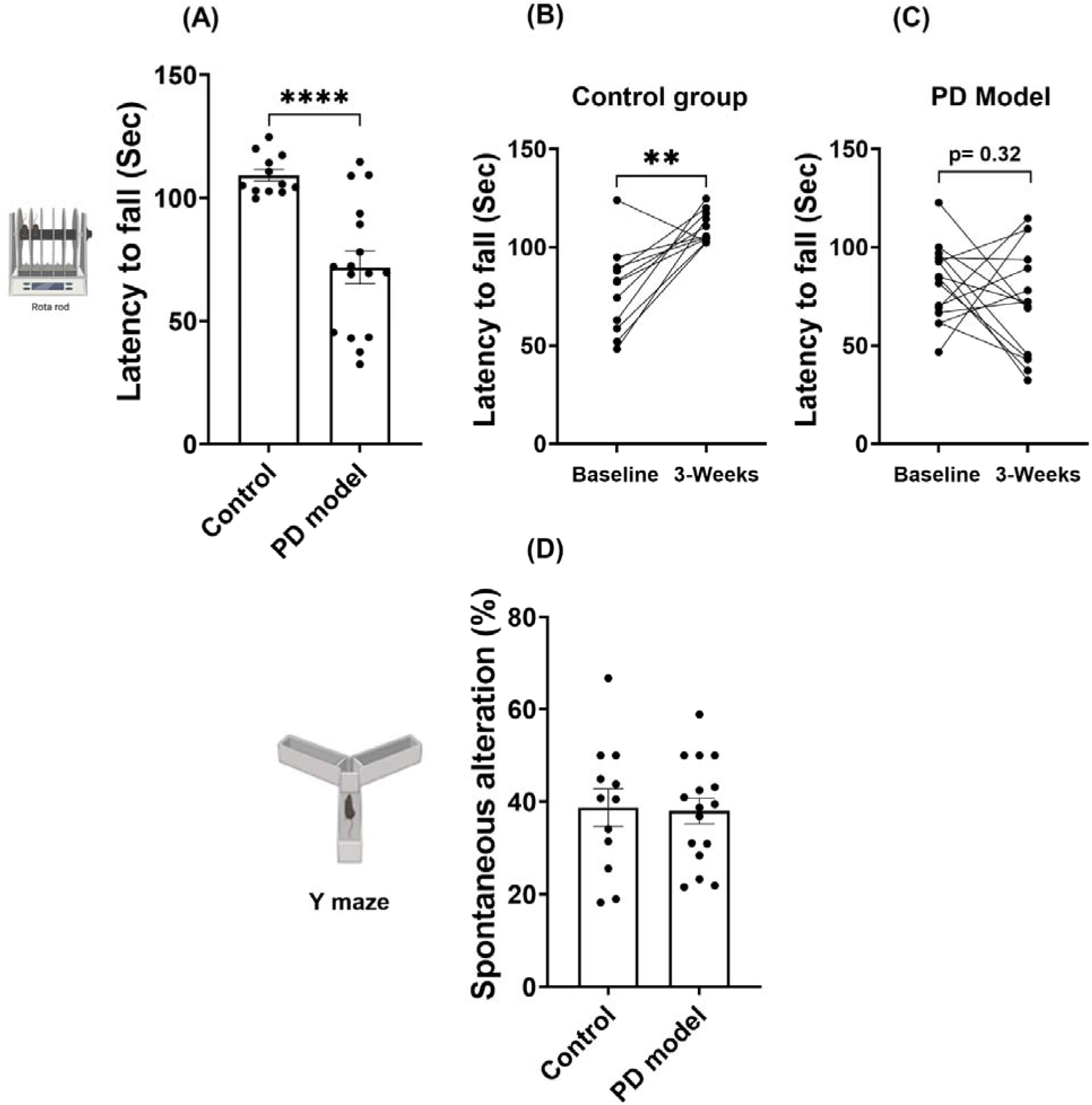
Intraperitoneal rotenone impairs motor function and motor learning at the acute phase. (A) Latency to fall during the rotarod task after 3 weeks of rotenone exposure. (B-C) Performance changes between baseline and after treatment completion in control mice (B) and PD model (C). (D) Spatial working memory alteration in Y-maze tests. Data are shown as mean ± SEM, two-tailed Welch’s *t* test (for A and D) and two-tailed paired *t*-test (for B and C). Control *n* = 12, PD model *n* = 16; *p < 0.01 (**), ****p < 0.0001*.

### Cellular changes in TH^+^ levels, dopaminergic cell density, and neuroinflammation during the acute phase

SNc is a histologically continuous structure composed of genetically and functionally diverse dopaminergic neurons in the middle brain^25–28^, and PD patients show prominent lesions in this region. We first examined dopaminergic lesions during the acute phase. Rotenone induced a strong reduction of TH fluorescence in the SNc (Figure 4A-C), particularly in the middle ventral SNc (Figure 4B), known to express SOX6+/Aldha1+ (but lack Otx2) ^26,27^. TH^+^ cell number and density in SNc decreased by ∼40% in rotenone-treated mice as compared with controls (Figure 4E-F). Interestingly, unlike SNc, striatum TH fluorescence was not changed, even in the dorsal lateral striatum that receives prominent SNc axonal projection^29^ (Suppl. Figure 1A-C). High-resolution imaging at this subregion confirmed the presence of intact TH^+^ axons and their varicosities. This is consistent with variable findings in striatal DA fibers^30^ and their compensatory sprouting in neonatal mice^31^, but contrasts with striatal DA axon loss and hypertrophy in rats^15^ with delayed SNc lesion in 6-OHDA model^32,33^ and MPTP model^34–36^, implying a different mechanism of toxicity in these PD models.

**Figure 4.**
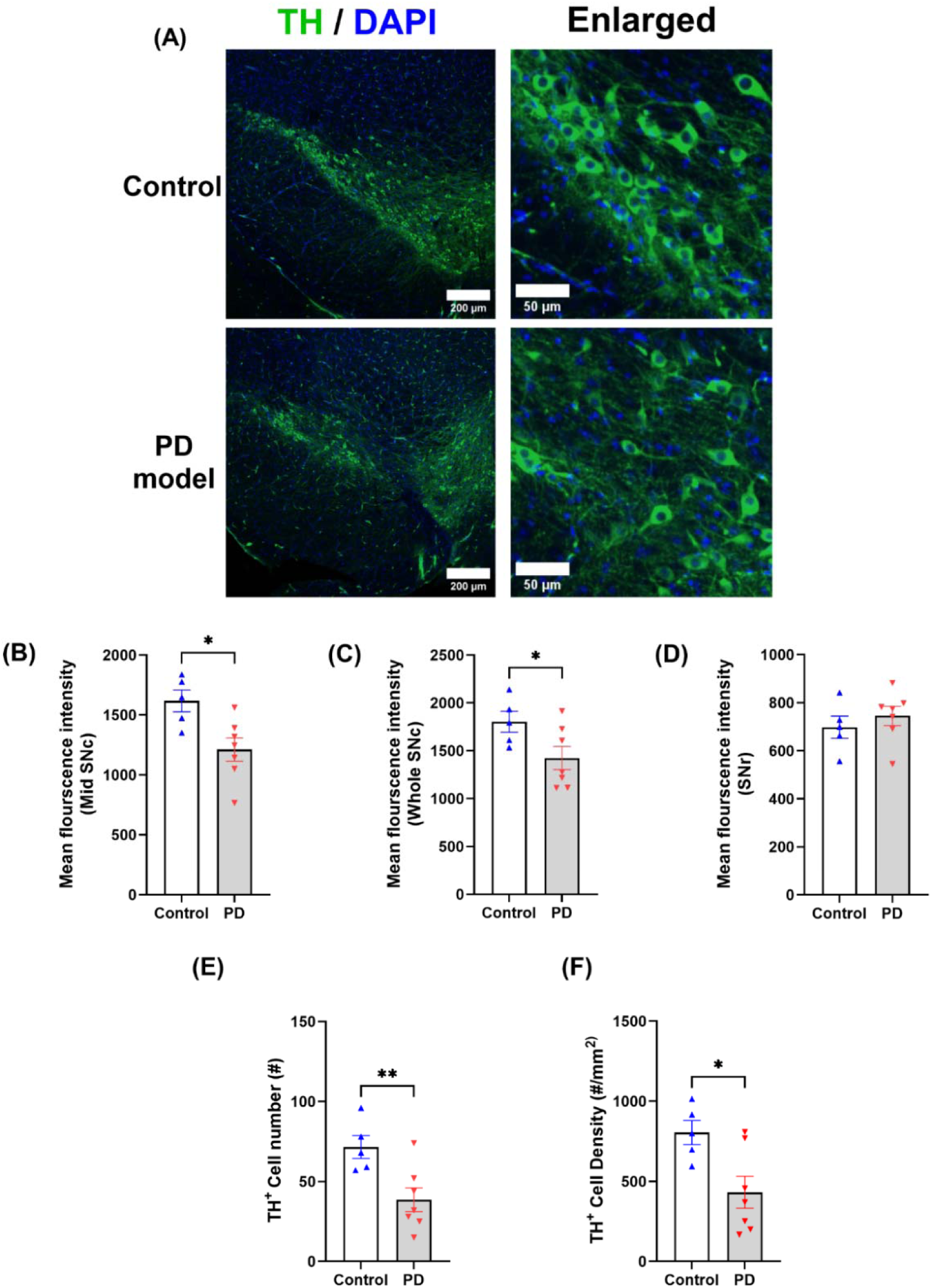
Chronic rotenone exposure causes dopaminergic neuron degeneration in SNc at the acute phase. (A) Representative confocal images of tyrosine hydroxylase (TH, green) with nuclear counterstain (DAPI, blue) in control and rotenone-treated mice. (B-D) Mean fluorescence intensity (MFI) of TH in the middle region of SNc (B), the entire SNc region (C), and SNr (D). (E) The number of TH^+^ neurons in the SNc. (F) TH^+^ cell density in the SNc. TH^+^ neurons were counted when TH^+^ spots co-stained with DAPI, and ROIs were drawn manually based on TH staining and local anatomy in Fiji (See Methods). Statistical analyses were performed using two-tailed Welch’s *t*-tests. Control *n* = 5, PD model *n* = 7; * *p < 0.05,* ** *p* < 0.01.

Next, we examined microglia changes in the middle brain. The purinergic P2RY12 receptor (P2RY12) is exclusively expressed on microglia and required for microglia migration and response. Its levels reduce upon microglia activation and can be lost upon microglia dystrophy ^37,38^. Inhibiting P2RY12 attenuates neurodegeneration in the MPTP-induced PD model ^39^. We used P2RY12 to probe neuroinflammation in our rotenone model and found no apparent changes in P2RY12 fluorescence intensity and microglia density (Supp. Figure 2). The absence of P2RY12 changes implies a limited role of neuroinflammation in this model, which is unexpected since other PD models induced by lipopolysaccharide (LPS) ^40^ or α-synuclein fibril injection ^41^ are accompanied by microglia activation.

### Locomotion deficits and anxiety behaviors are resolved by 12 months

After establishing the robust Parkinsonism at the acute phase, we examined their long-term recovery at 12 months. To our surprise, most of the deficits observed in the acute phase were resolved, and the rotenone-treated mice behaved normally as controls. As shown in the open field tests, rotenone-treated mice showed similar locomotion as controls (Figure 5). Both groups exhibited comparable outcomes in travel distance, average speed, and mobility time. Anxiety-like behavior, as measured by inner zone distance and outer zone distance in rotenone-treated mice were equal to that of controls (Figure 5D-F). The longitudinal results within 12 months revealed a full picture of rotenone toxicity in short and long term (Figure 5K). While locomotion was strongly impaired in the acute phase, it fully recovered to the control levels within ∼4 months.

**Figure 5.**
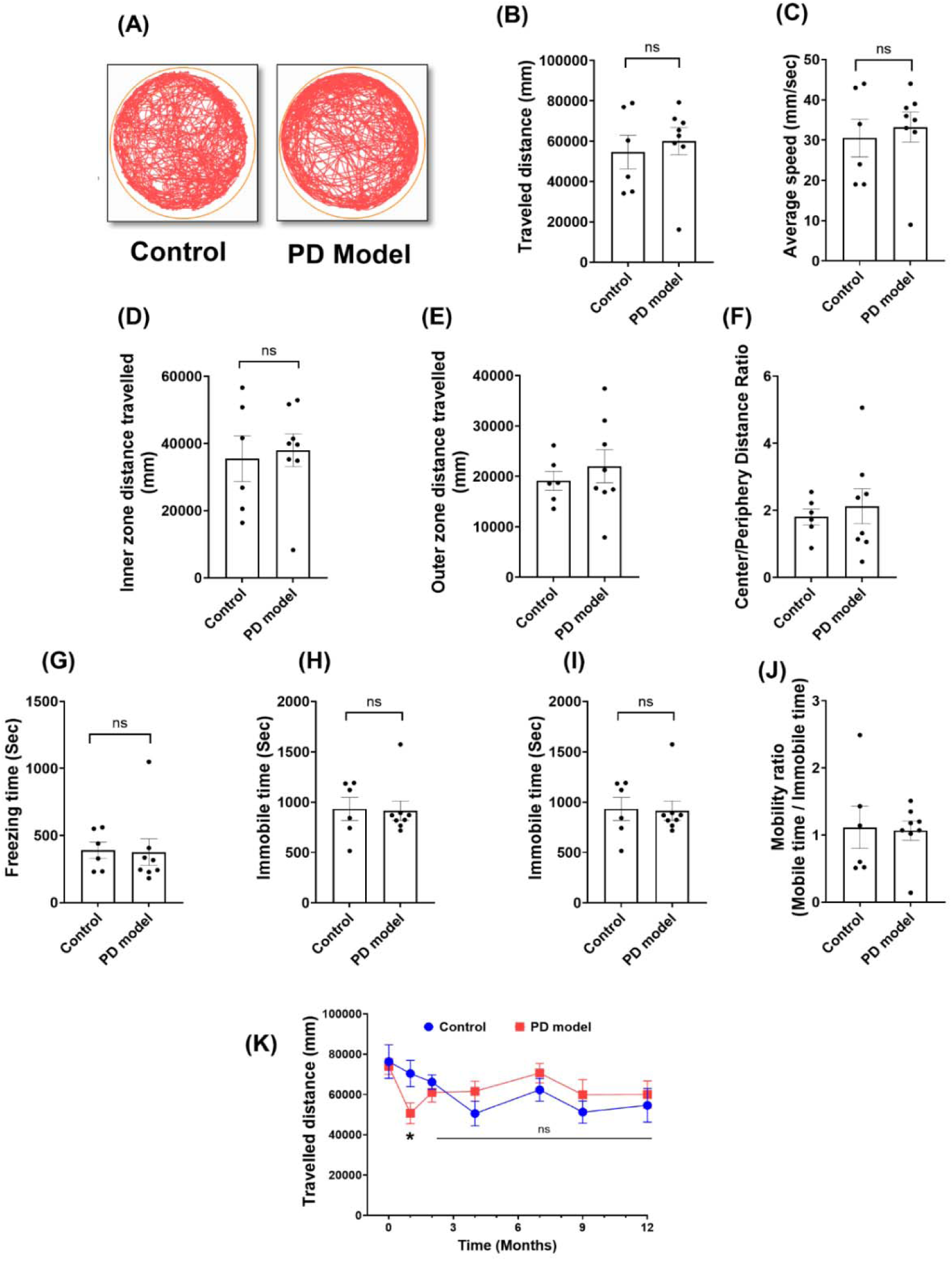
Long-term recovery of locomotor function at 12 months following 3 weeks of rotenone exposure. (A) Representative open-field movement tracks from control and rotenone-treated mice. (B-J) locomotor activity showing the changes in total distance traveled (B), speed (C), center zone distance (D), outer-zone distance traveled (E), center-to-periphery distance ratio (F), freezing time (G), mobile time (H), immobile time (I), and mobility ratio (J). (K) Longitudinal analysis of the total distance traveled at different times. The first point in each group represents the baseline before treatment, plotted as the “0 month” point in (K). Data were presented as mean ± SEM, Two-tailed Welch’s *t* tests. Sample size: Baseline and 1 month Control *n* = 12, PD model *n* = 16; 1 to 7 months Control *n* = 6, PD model *n* = 10; 9 to 12 months: Control *n* = 6, PD model *n* = 8. *P < 0.05,* ** *p < 0.01*.

EPM tests at 12 months further showed no difference between the two groups (Figure 6). They showed similar results in total travel distance, open arm entry, open arm time and distance. This is consistent with the results in the open field tests showing indistinguishable inner/outer distance (Figure 5D-F). As expected, the Y-maze tests showed no changes between the two groups in the acute phase.

**Figure 6.**
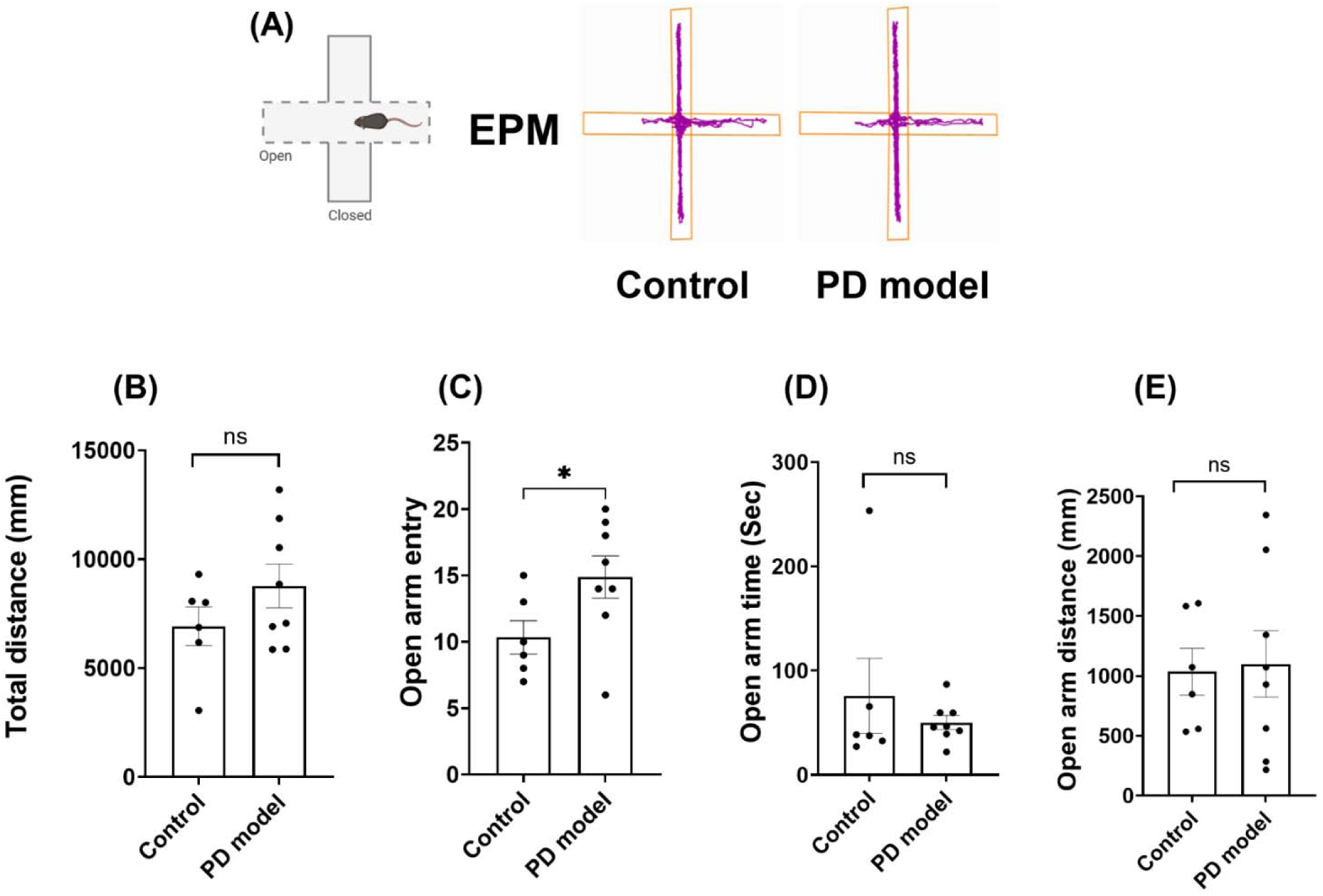
Long-term recovery of anxiety-like phenotypes at 12 months. (A) Representative mouse trajectories in the EPM at 12-months. (B-E) Quantification analysis for the total distance traveled (B), open arm entries (C), open arm time (D), or open arm distance (E). Data are shown as mean ± SEM; two-tailed Welch’s *t* tests; control *n* = 6, PD model *n* = 8 mice; ns, not significant.

These results indicate that rotenone-induced Parkinsonism, including impaired locomotion and non-motor function, in the acute phase are fully reversible in the long-term. These findings uncover the previously unappreciated capacity of the dopaminergic system in its resilience and adaptability.

### Impaired motor balance and coordination recover by 12 months

Next, we measure the long-term recovery of motor balance and coordination using rotarod assays. Rotenone-treated mice performed the task equally well as control mice at 12 months (Figure 7A), suggesting the recovery of motor balance and coordination. The longitudinal analysis of motor function revealed dynamic changes in the same mice within 12 months (Figure 7B). Rotenone treatment significantly worsened the performance of the rotarod task in the acute phase (after the 3^rd^ week), but the rotenone-treated mice became indistinguishable from the age-matched controls at 7 months. Notably, both groups exhibited a decline in motor performance over the course of the experiments. The transient increase of the control group at the 1^st^ month before performance decline may arise from the initial motor learning, which was impaired in rotenone treated group. The curve divergence of the two groups at the acute phase speaks for the effectiveness of rotenone treatment, and their convergence at the later stage supports the long-term recovery of rotenone-induced damage.

**Figure 7.**
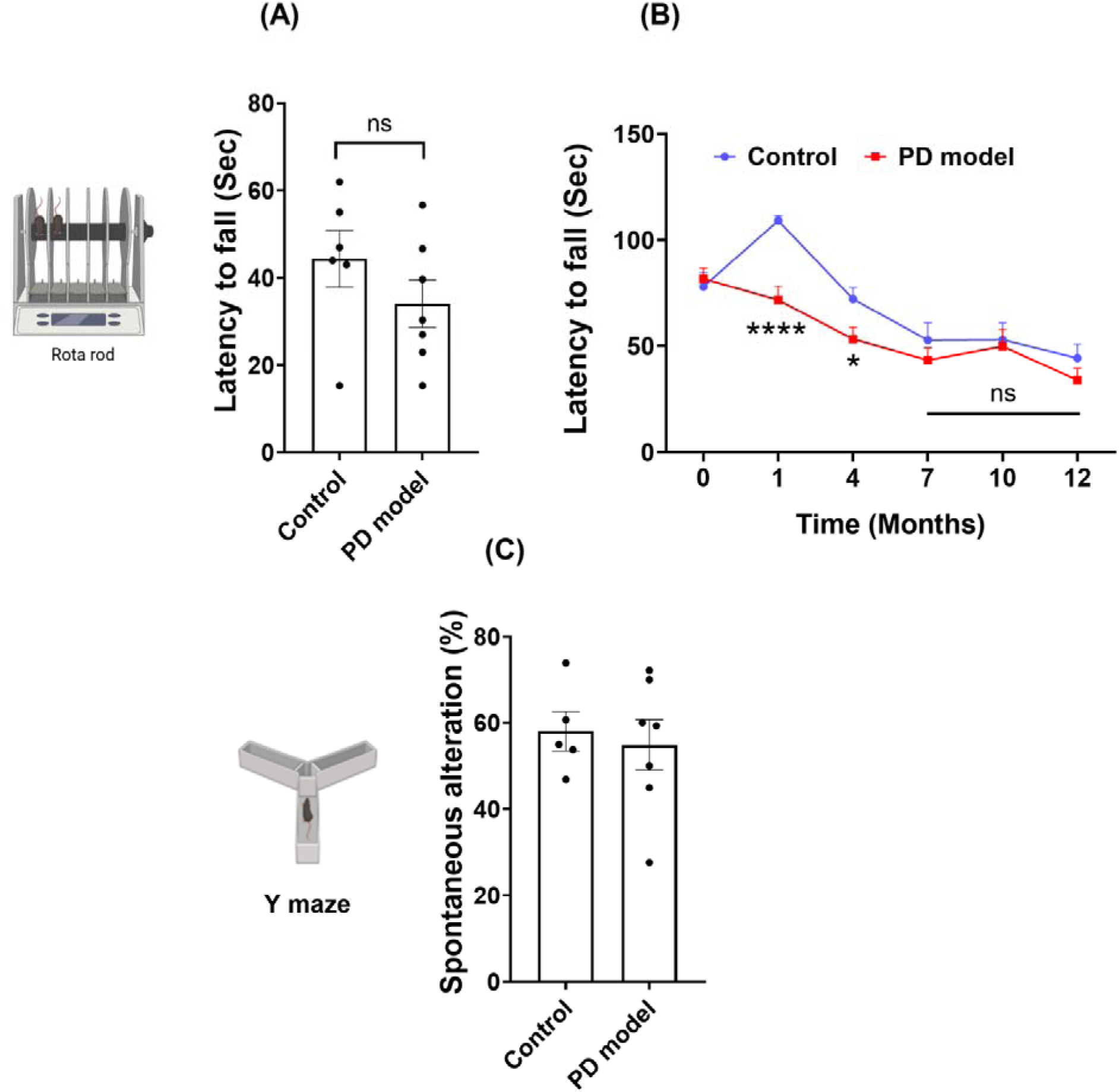
Long-term recovery of motor dysfunction at 12 months. (A) Rotarod task performance was not significantly different between the PD model and controls at 12 months. (B) Longitudinal analysis of rotarod task performance at different times. Note: the significantly worse performance in the PD mice at 1 and 4 months, but similar performance at 7 months and later. (C) Y-maze tests at 12 Months. Data are shown as mean ± SEM; two-tailed Welch’s *t* test. Sample size: Baseline to 1 month: Control *n* = 12, PD model *n* = 16. 12-Months-Rotarod, control *n* = 6, PD model *n* = 7; Y maze, control *n* = 5, PD model *n* = 7; *\* p < 0.05, **** p < 0.0001*; ns = not significant.

### Cellular pathology recovers after 12 months

Given the notion that adult brain neurons rarely regenerate and neurodegeneration is irreversible, the observed long-term recovery to control levels is surprising. To better understand its mechanism, we examined middle brain cellular changes at 12 months. Figure 8 shows TH fluorescence levels and TH^+^ cell density in the SNc between the two groups. In comparison, rotenone-treated mice still had slightly lower levels of TH fluorescence than the age-matched controls (Figure 8A-C). Notably, TH^+^ cell density in control SNc reduced significantly at 12 months (Figure 8F) as compared with at 1 month (Figure 4F). This is consistent with age-dependent decline (>30%) of SNc TH^+^ neuron numbers reported in rats^42^ and humans ^43,44^. However, this age-dependent decline of TH^+^ cell density in the rotenone-treated group was much more limited (Figure 4F and 8F). Additionally, TH levels from the dorsal lateral striatum remained similar between the two groups (Suppl. Figure 1D-F).

**Figure 8.**
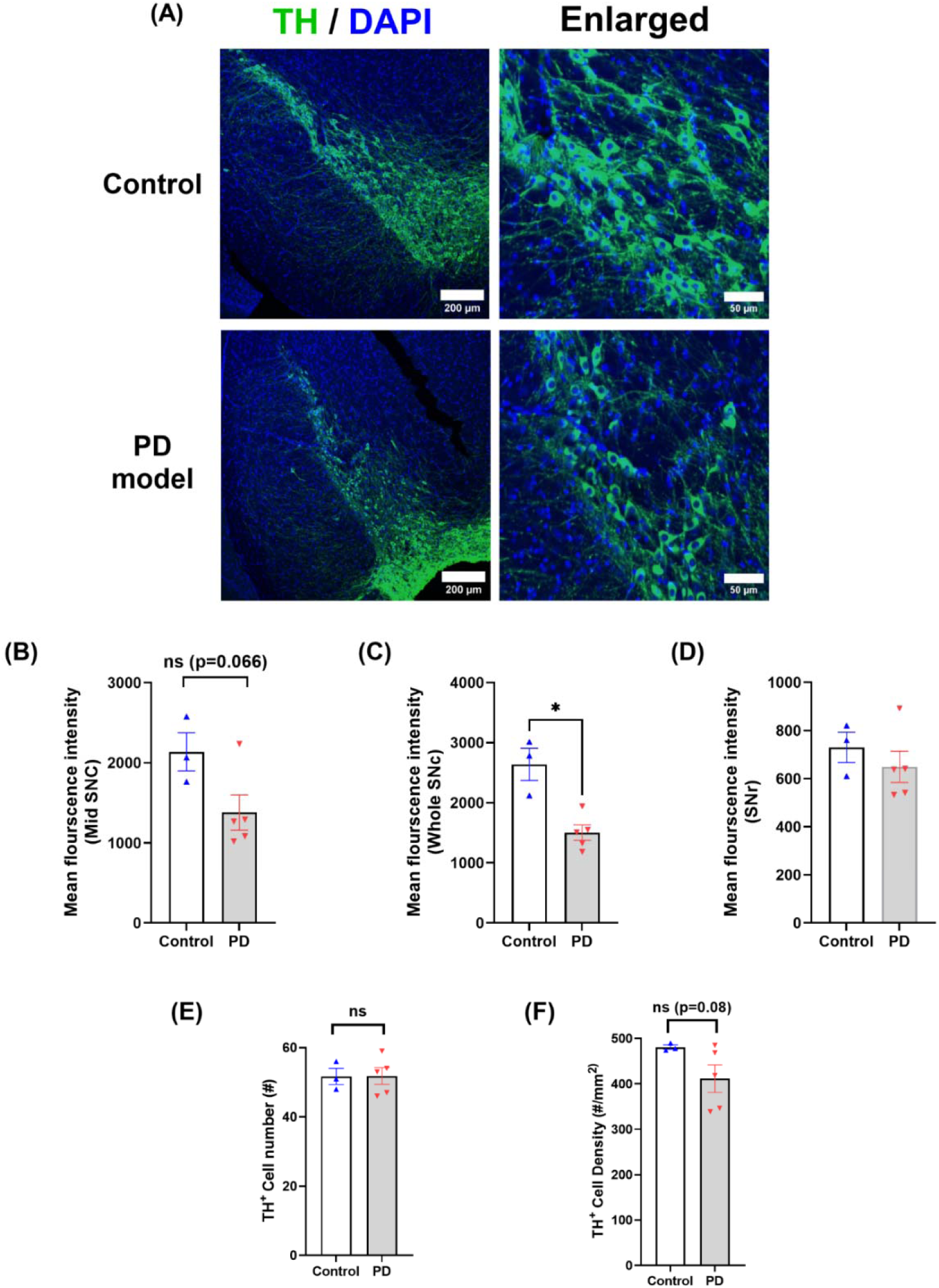
Long-term changes of dopaminergic neurons in SNc at 12 months. **(A)** Representative confocal images of middle brain sections stained with TH (green) and nuclei (DAPI, blue) from control and rotenone-treated groups at 12-months. (B-D) Quantification of TH levels in the mid-SNc (B), the entire SNc (C), and SNr regions (D). (E-F) Quantification of TH^+^ cell counts (E) and TH^+^ cell density (F) in the SNc region. (Mean ± SEM, two-tailed Welch’s t test; Control *n* = 3, PD model *n* = 5; *\* p < 0.05;* ns = not significant.

We further examined P2RY12 to see if there was delayed microglia activation. SNc imaging showed slightly higher P2RY12+ levels in SNc than in SNr, but this was comparable between the two groups of mice (Supp. Figure 3A-E). However, SNr P2RY12 cell number and density were both lower in rotenone-treated mice than in controls (Suppl. Figure 3F-G), suggesting partial microglia activation in the SNr region. Dorsal striatal imaging revealed similar P2RY12 levels and microglia density between the two groups (Supp. Figure 3H-J).

Collectively, these data suggest that at 12 months, SNc TH levels remained lower, but TH+ cell density in the rotenone-treated group was recovered to the levels of age-matched controls.

## Discussion

This work focuses on the long-term recovery of rotenone-induced Parkinsonism. We first established the robust PD phenotypes in the acute phase and then monitored their longitudinal changes for 12 months. The results uncover a surprising reversibility of the impaired motor and non-motor function, which has direct implications for understanding the long-term compensation and plasticity of the dopaminergic system in PD.

Rotenone models have been increasingly used to mimic progressive features of PD^7^ since its first demonstration in Sprague-Dawley and Lewis rats^6^. While rat rotenone models have been well studied ^7,15,45^, mouse rotenone models have been less explored, with mixed results reported^9^. The oral rotenone mouse model was used to study PD ^14,21,46^ and to evaluate therapeutics^47–49^, but its effectiveness as a PD model was challenged recently^24^. Other routes of exposure, such as subcutaneous administration at low doses (2.5-4 mg/kg/day) ^50^, intraperitoneal (1-10 mg/kg/day for 3 weeks)^51^, or environmental exposure via breeding cages (5 mg/kg/day for 2–6 weeks) ^20^, have also been explored for modeling PD. In this work, we demonstrated that rotenone intraperitoneal injection induced robust Parkinsonism, which occurred in the acute exposure phase. Chronic rotenone IP injection gradually induced PD-like deficits, including hypokinesia, impaired motor coordination and motor learning, as well as non-motor dysfunction such as increased anxiety. Along with these functional deficits, the cellular pathology becomes prominent in parallel, including dopaminergic neuron loss in SNc and reduced TH expression. Interestingly, these changes were confined to SNc, without changes in the striatum. This may be partially attributed to multiple origins of TH^+^ axons at the striatum or compensatory axon fiber sprouting ^31^. Notably, our IP model displayed a much higher survival rate, and all the mice survived except a few mice were euthanized for non-specific conditions. This contrasts with the 85% mortality within 56 days of oral rotenone exposure previously reported^14^, implying lower non-specific toxicity of rotenone in this PD model.

Most rodent studies focused on the rotenone effect at the acute phase, leaving its long-term consequences unclear. Here, we performed a longitudinal study for 12 months after rotenone exposure. We found that the parkinsonism developed in the acute phases, such as locomotion deficit, anxiety, and motor balance and coordination, were fully reversible in the long term. These results provide the first direct evidence on the adaptation and resilience in PD-associated neural circuits. Although the mechanisms remain unclear, it may involve compensatory neural plasticity or circuitry remodeling linked with motor and non-motor deficits, and these mechanisms may be exploited to gain insights into how to slow PD progression. This is the first study in mice that tracks PD onset, development, and reversal for over a year. It revealed an age-dependent decline of motor performance in both control and PD groups. The initial performance increase in the control but not in the rotenone group may be attributed to different motor learning. While the curve divergence between two groups speaks for the effect of rotenone treatment at the acute phase, the curve convergence at the later stage illustrates the long-term recovery of deficits in vivo.

The long-term recovery of DA neuron density is surprising since neurodegeneration was thought to be irreversible, and most adult brain neurons cannot regenerate^52,53^. On the other hand, it becomes increasingly clear that the dentate gyrus and subventricular zone retain the ability of adult neurogenesis under physiological conditions ^52,54^. Further, certain pathological conditions (e.g., brain ischemic injury) can activate neurogenesis in the hippocampus ^53,55–57^. The SNc region contains heterogeneous dopaminergic neurons^25,27,28^, with highly diverse genetic profiles. Chronic rotenone exposure may reprogram certain dopaminergic neurons to regain regeneration ability. In addition, recent work reported that both adult serotonin^58^ and norepinephrine neurons^59^ possess the intrinsic ability to regrow their axons upon injury^60^. Dopaminergic axons share some similarities in the long-range, extensive branching and spreading projection with modulatory roles, and thus their axons in the striatum may regain a similar ability for axon regrowth or compensatory re-sprouting upon rotenone-induced stress. Indeed, neonatal SNc lesions induced by 6-OHDA unilateral injection have substantial capacity for compensatory sprouting of DA axons^31^, leading to less TH reduction in the striatum than in SNc. Lastly, it needs to keep in mind the species-specific differences between mice and humans. Mice have much shorter lifespans and smaller cell sizes, and they also exhibit critical molecular differences in the PD context, such as different dopamine metabolism^61^, lack of neuromelanin, divergent properties in α-synuclein and LRRK2^62^, and glia gene profiles^63–65^. Thus, future work is needed to verify whether these results can be extrapolated to humans.

The progressive development of motor deficit and loss of dopaminergic neurons at the acute phase supports a critical role of mitochondrial dysfunction in PD pathogenesis, which is aligned with the positive correlation between pesticide exposure history and PD risk. However, the recovery of these deficits after rotenone withdrawal uncovers a limited long-term consequence of rotenone exposure, questioning the long-held notion of pesticide history as a cause of PD. Indeed, while epidemiological studies support a generic correlation between pesticide exposure history and PD incidence, numerous methodological limitations prevent a definite conclusion on the causal role of the exposure^3,4^.

Together, this work has established the effectiveness of rotenone exposure to induce Parkinsonism and further evaluated the long-term effects of rotenone exposure. The longitudinal data reveal the reversibility of PD-like deficits in the long-term, uncovering a previously underappreciated capacity of resilience and adaptation of the dopaminergic system in vivo. This property may be exploited further for preventing PD progression.

## Methods

### Animals and Housing

All experiments were performed on age-matched male C57BL/6J mice (Jackson Laboratory, USA). Mice were housed in a pathogen-free, temperature-controlled (22 ± 2 °C) and humidity-controlled (45–60%) facility under a regular 12 h light/dark cycle, with standard polycarbonate cages in the cages. All cages were supplied with paper bedding and nesting materials. Food (standard rodent chow; LabDiet 5001) and water were available ad libitum. Mice showing severe weight loss (>20% baseline) or distress were excluded per institutional animal welfare protocols. All procedures were approved by the Institutional Animal Care and Use Committee (IACUC) of the Medical College of Wisconsin and followed the NIH Guide for the Care and Use of Laboratory Animals.

Male mice at 2 months of age were used for experiments since males produced consistent results and were less variable than females^9,22,23^. Mice were acclimated to the behavioral core facility for at least 2 weeks to minimize environmental impact. To further reduce non-specific behavioral perturbation, cohorts were randomized into two groups with intraperitoneal (IP) rotenone as the PD model and vehicle as controls, housed together throughout the study, and tested side-by-side in parallel and by the same well-trained experimenter for all animals.

### Rotenone Preparation and Dosing

Rotenone (Cat#R8875, Sigma-Aldrich, USA) was administered to C57BL/6 mice via intraperitoneal injection. Briefly, rotenone powder was dissolved in 100% dimethyl sulfoxide (DMSO) to generate a 50× stock solution, aliquoted into amber septa vials to avoid light exposure, and stored at −20 °C. The stock solution was diluted in the medium-chain triglyceride Miglyol 812 (Code# 207878B, Medisca) to produce a final working solution of 2.5 mg/kg bodyweigh^15^ at a dosing volume of 10 µL/g body weight (98% Miglyol + 2% DMSO). The mixture was prepared fresh every other day. IP injections were administered once daily for 21 consecutive days. Control mice received equivalent volumes of vehicle. Throughout treatment, mice were monitored daily for body weight, grooming, posture, mobility, and general health, with any adverse signs recorded and animals supported per institutional welfare guidelines.

### Evaluation of motor and non-motor function

Open Field Test: Spontaneous locomotor activity was assessed in the open field tests (OFT). We used an apparatus consisting of four large circular arenas (diameter: 19 in, height: 12.5 in; corrugated baseboards). Before testing, animals were habituated to the behavioral suite for 30 minutes. Mice were individually placed in the center of the arena and allowed to explore freely for 30 minutes. Between each trial, arenas were thoroughly cleaned with 70% ethanol to remove olfactory cues. All experiments were conducted under standard lighting conditions with continuous white noise to minimize external disturbances. Mouse behavior was recorded by an overhead camera, analyzed using ANY-maze software (Stoelting Co., USA) for quantification of total distance traveled, average speed, time spent in the inner zones and outer zones, mobility, immobility, and freezing duration. Mice were assessed right before treatment (as the baseline) and then tested weekly to monitor deficit onset and progression, with the results measured shortly after the 3-week treatment as the acute phase. After that, mice were either used for cellular pathology characterization or kept for evaluating longitudinal recovery from 2 to 12 months and pathological examination at the end of the study.

Elevated Plus Maze (EPM) Test. Anxiety-like behavior was assessed using the elevated plus maze (EPM; San Diego Instruments, USA; Product #7001-0316). The apparatus consists of two open arms (length: 12 in; width: 2 in; wall height: 15.25 in) and two closed arms (length: 26 in; width: 2 in; wall height: 21.25 in) arranged at right angles around a central platform. Mice were habituated to the behavioral suite for 30 minutes before testing, and tests were performed under standard lighting with continuous white noise. Each mouse was placed in the central platform facing an open arm and allowed to freely explore for 5 minutes. The maze was thoroughly cleaned with 70% ethanol between trials to eliminate olfactory cues. Behavior was recorded using an overhead camera and analyzed by ANY-maze software (Stoelting Co., USA) for automated scoring. Results were quantified using the total distance traveled in the maze, the number of open arm entries, time spent in the open arm, and the distance traveled in the open arm.

Rotarod Test. Motor coordination and balance were assessed using a Rotarod treadmill (IITC Life Sciences, USA; Product #755) equipped with rat drums (diameter: 3.75 in; lane width: 4 in; lane height from drum to base: 12 in). Before testing, animals were habituated to the behavioral suite for 30 minutes. Mice were placed individually on the rotating drum, and latency to fall was automatically recorded by the system. Testing was performed under an accelerating paradigm, with a start rotation speed of 4 rpm and continuously increased to 40 rpm over a 5-minute trial. Each mouse underwent three trials per day, separated by 30-minute inter-trial intervals, for three consecutive days. The third trial was used to analyze the final performance. Between each trial, the instrument was thoroughly cleaned with 70% ethanol to remove olfactory cues. All experiments were conducted under standard lighting conditions with continuous white noise to minimize external disturbances. Mice were tested right before treatment (as the baseline) and then tested weekly to monitor the deficit onset and progression, with the results measured shortly after the 3-week treatment as the acute phase. Mice were tested further after completion of rotenone exposure (IP, 2.5 mg/kg/day × 21 days) longitudinally from 2 to 12-months post-treatment. Latency to fall was averaged across trials for each mouse.

Y-Maze Test. Working memory was evaluated using the spontaneous alternation Y-maze paradigm (San Diego Instruments, USA; Part #7001-0424 Y). The maze was constructed of 0.25″ acrylic with clear walls and beige floors, consisting of three arms (length: 15 in; width: 3 in; wall height: 5 in) arranged at 120° angles. Mice were habituated to the behavioral suite for 30 minutes prior to testing, which was performed under standard lighting with continuous white noise to minimize external disturbances. Each mouse was placed individually at the end of one arm and allowed to freely explore all three arms for 10 minutes. Arm entries were recorded using an overhead camera system, and spontaneous alternation behavior was calculated as the number of sequential visits (ABC, ACB, BAC, BCA, CAB, CBA) to three different arms divided by the total possible alternations, expressed as a percentage. Between trials, the maze was thoroughly cleaned with 70% ethanol to remove olfactory cues.

### Brain Tissue Preparation, Immunofluorescence Imaging and Analysis

Brain tissues were prepared as previously reported^66,67^, with minor modifications. Briefly, mice were euthanized by rapid decapitation at the middle of the 4^th^ week of rotenone exposure, and brains were immediately isolated and immersed in ice-cold phosphate-buffered saline (PBS). Whole brains were immersed in freshly prepared 4% paraformaldehyde (PFA) and 4% sucrose in PBS for 24 h at 4 °C with gentle shaking. Following fixation, brains were transferred to 30% sucrose in PBS and stored at 4 °C until sectioning. Coronal sections (30 µm) were cut using a vibratome, with the region of interest identified visually and under a microscope and mounted on glass slides. The tissue sections encompassing substantia nigra pars compacta (SNc), substantia nigra pars reticulata (SNr), and striatum (ST) were identified by their cellular architecture and adjacent brain structures in the middle brain. SNc is an irregular-shaped nucleus concentrated with dopaminergic neurons, spanning a narrow, elongated region up to ∼1.4 x1.4 x 1 mm at medial-lateral, rostral-caudal, and dorsal-ventral directions (Allen Mouse Brain Atlas). In the coronal section of the middle brain, it displays as a wing-like structure, with dorsal SNc gradually merged with lateral Parabrachial pigmented nucleus (PBP), a part of the ventral tegmental area (VTA) toward the midline. To minimize the subregional effect of slices, tissue sections from similar SNc anatomic levels were used for comparison across individual mice and groups.

For immunofluorescence staining, brain sections were washed in PBS and blocked for 1 h at room temperature in PBS containing 0.4% Triton X-100 and 5% normal goat serum. Sections were incubated overnight at 4 °C with primary antibodies diluted in blocking buffer. The next day, sections were washed and incubated for 2 h at room temperature with secondary antibodies conjugated with Alexa Fluor dyes or CF-dyes (1:300, Invitrogen). Nuclei were counterstained with DAPI (1 µg/mL), and sections were mounted with ProLong Gold antifade mounting medium (Thermo Fisher). Primary antibody used here: rabbit anti-tyrosine hydroxylase (TH; 1:400, Millipore #AB152; 1:400, Immunostar Cat# 22941), P2RY12 (Cat#848002, 1:50, Biolegend).

Confocal imaging was performed primarily using an Andor BC-43 spinning disk confocal microscope (Oxford Instruments, UK) equipped with an sCMOS camera and objectives of 10×, 20×, and 60× oil. Some experiments were also tested with a Nikon spinning disk confocal microscope, as described previously ^68–70^, and an Abberior STED microscope. For each batch of experiments, all images were acquired using identical acquisition parameters (laser intensity, exposure time, and camera binning) to ensure consistency across samples. Images for comparison were presented with the same display settings.

Quantitative analysis was performed on 2D images, and regions of interest (ROIs) in the SNc, SNr, and striatum were delineated using the free-hand selection tool in ImageJ and Fiji. The ROIs of SNc were defined based on TH staining pattern and local neural anatomy, excluding the PBP and VTA complex. Striatum images were acquired from the comparable sites/areas at the dorsal lateral striatum from each coronal section since this region is a major site of axonal projection from SNc dopaminergic neurons. Mean fluorescence intensity, percentage area of positive signal (% area), and cell density were measured within the ROIs, and the results were from different batches of experiments. *TH^+^ neurons were visually identified by typical TH fluorescence spots diffusing around a nucleus after image zoom-in*.

### Statistical Analysis

All data were presented as mean ± standard error of the mean (SEM), with individual data points representing biological replicates or individual mice. Statistical analyses were performed using GraphPad Prism (version 10.6.0; GraphPad Software, USA). Comparisons between two groups were conducted using a two-tailed Student’s *t*-test with Welch’s correction (Welch’s *t*-test) to account for unequal variances. For multi-group comparisons (e.g., column graphs), one-way or two-way analysis of variance (ANOVA) was applied, followed by Tukey’s multiple-comparisons post hoc test. For longitudinal or repeated time-point analyses (e.g., time-plot graphs), two-way ANOVA with Sidak’s multiple-comparisons post hoc test was used. Statistical significance was defined as \**p* < 0.05, \*\**p* < 0.01, \*\*\**p* < 0.005, \*\*\*\**p* < 0.001.

## Acknowledgement

This work is supported partially by Awards from the National Institutes of Health (NIH, R01DK132088, R01DK133326, and R01AG079257 to X.L.), the Medical College of Wisconsin (to X.L.), and Advancing a Healthier Wisconsin Endowment (AHW) (FP00028084 to P.J.). P.J. is partially supported by the AHW Post-doc Research Seed Grant. We thank the Oxford Instruments Center for Advanced Microscopy and the EM core for imaging experiments and analysis. The FACILITY LINE STED microscope was purchased via an NIH high-end Instrument S10 Award (1S10OD034247-01A1 to X.L.). We thank other members of the Lou Lab for technical support and comments on the manuscript.

---

**Supplementary Figure 1.**
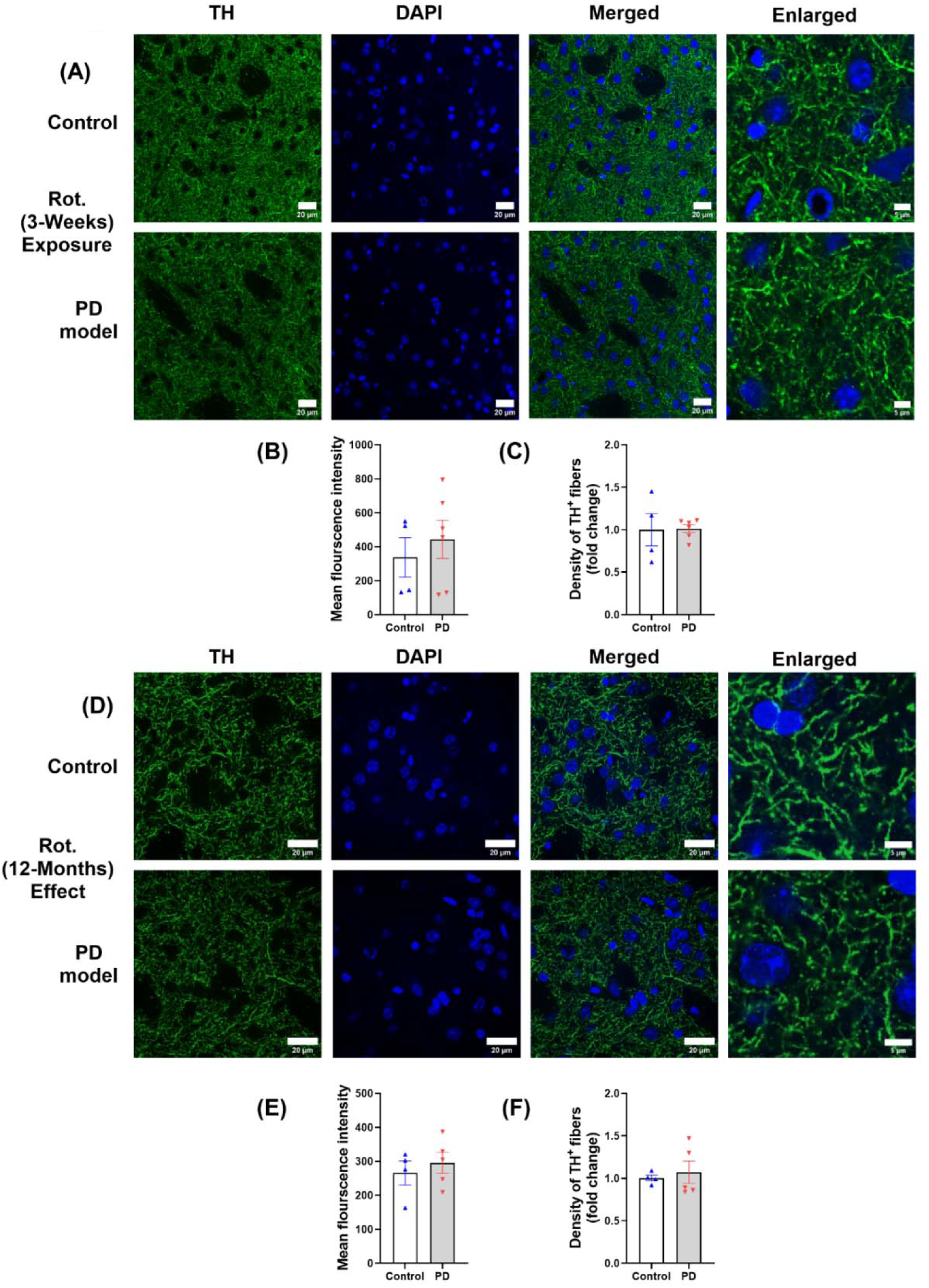
Striatal TH was preserved after both short and long-term rotenone exposure in mice. (A) Representative confocal images of TH immunofluorescence (green) and nuclear counterstain (DAPI, blue) in the striatum after 3 weeks of rotenone exposure. The far-right panel shows enlarged views of individual TH^+^ axons. (B-C) Intact TH fluorescence and density in the striatal dorsal lateral region at 1 month. (D-F) like (A-C) but at 12 months. Data are presented as mean ± SEM, Mann-Whitney nonparametric test; 3-weeks (control *n* = 4, PD model *n* = 6), 12-months (control *n* = 4, PD model *n* = 5).

**Supplementary Figure 2.**
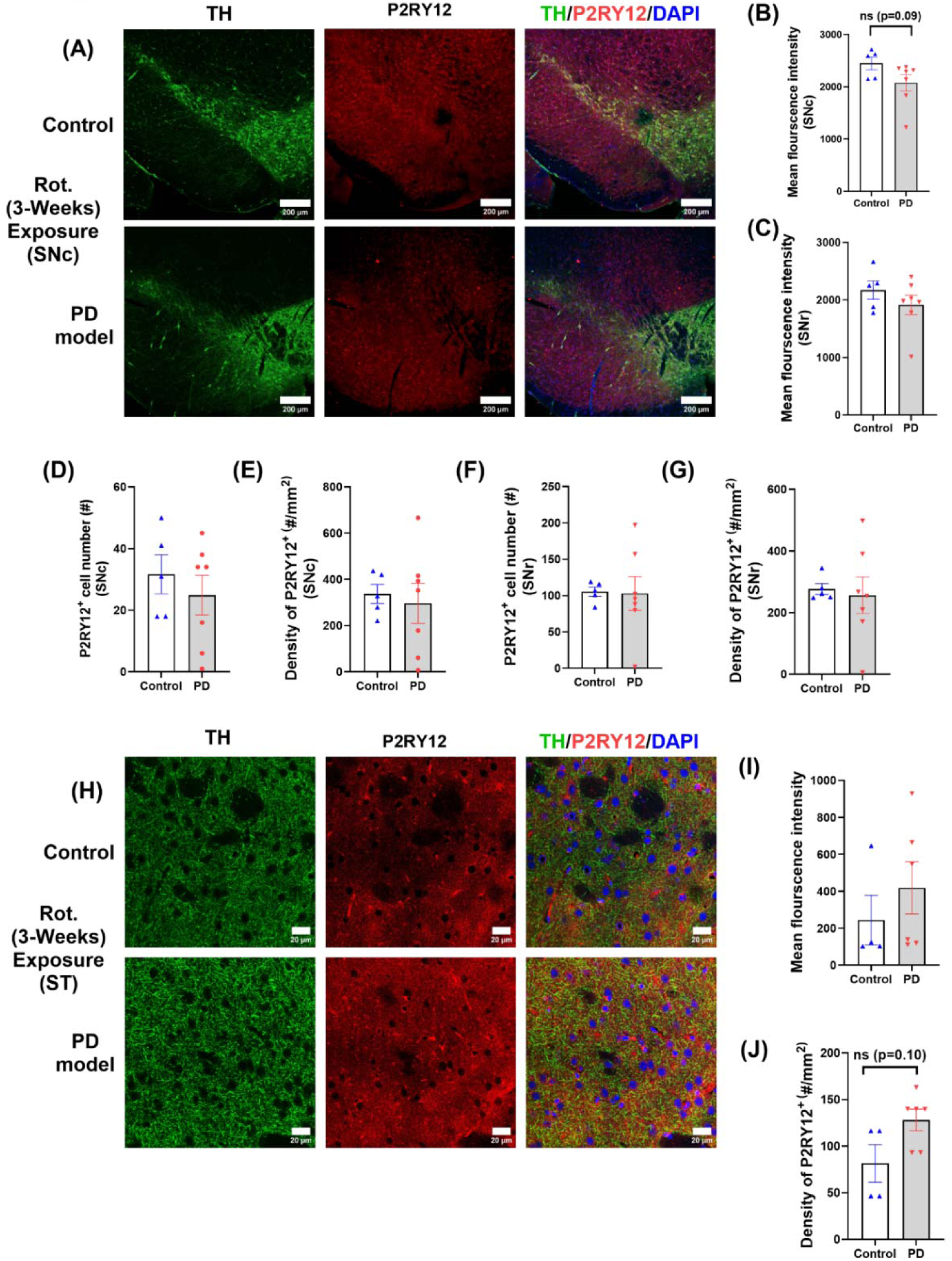
Characterization of microglial P2RY12 changes in SNc and striatum at the acute phase of rotenone exposure. **(A)** Representative confocal images of SNc showing TH (green), P2RY12 (red), and nuclei (DAPI; blue) from coronal sections of control and rotenone-treated mice after 3 weeks of rotenone treatment. (B-C) Microglial P2RY12 fluorescence intensity in the SNc and SNr. (D-E) P2RY12□ microglial cell number and cell density in the SNc, and (F-G) P2RY12□ microglial cell number and cell density in the SNr. (H) Confocal images of TH (green), P2RY12 (red), and nuclei (DAPI; blue) in the dorsal lateral striatum. (I-J) Intact P2RY12 fluorescence intensity and P2RY12□ cell density. Data are presented as mean ± SEM; Mann-Whitney nonparametric tests; SNC (control *n* = 5, PD model *n* = 7), striatum (control *n* = 4, PD model *n* = 6); ns, not significant.

**Supplementary Figure 3.**
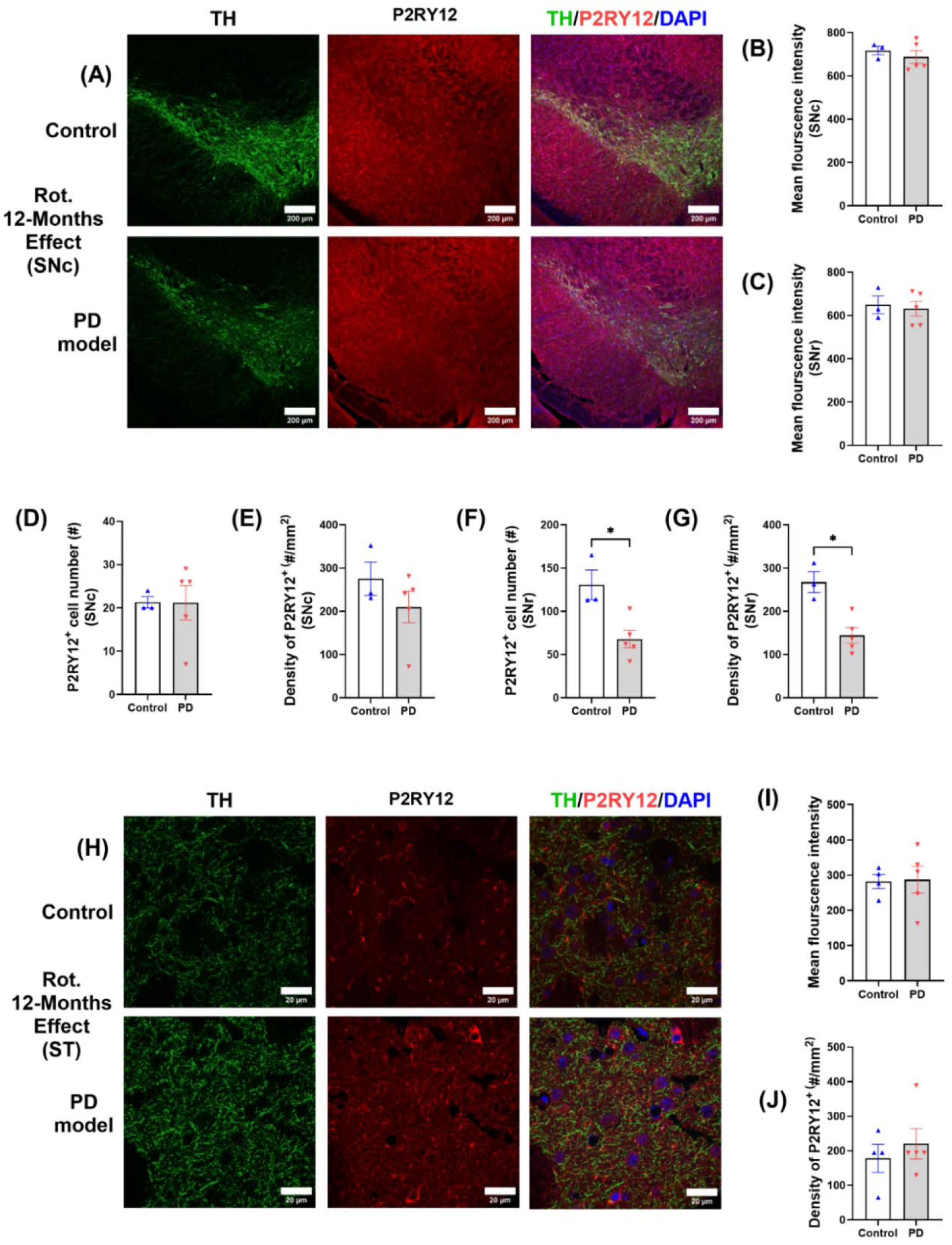
Microglial P2RY12 changes in the SNc and striatum at 12 months. **(A)** Representative confocal images of SNc regions showing TH (green), P2RY12 (red), and nuclei (DAPI; blue) from control and rotenone-treated mice at 12 months. (B-C) Microglial P2RY12 fluorescence intensity in the SNc and SNr. (D-E) P2RY12□ microglial cell number and cell density in the SNc. (F-G) □P2RY12 microglial cell number and cell density in the SNr. (H) Confocal images of TH (green), P2RY12 (red), and nuclei (DAPI; blue) in the dorsal lateral striatum at 12 months. (I-J) Intact P2RY12 fluorescence intensity and P2RY12□ cell density. Data are presented as mean ± SEM; two-tailed Welch’s *t* test; SNc (control *n* = 3, PD model *n* = 5), striatum (control *n* = 4, PD model *n* = 5); ns, not significant.

## References

1. Bloem, B.R., Okun, M.S., and Klein, C. (2021). Parkinson’s disease. The Lancet 397, 2284–2303.

2. Ye, H., Robak, L.A., Yu, M., Cykowski, M., and Shulman, J.M. (2023). Genetics and pathogenesis of Parkinson’s syndrome. Annual Review of Pathology: Mechanisms of Disease 18, 95–121.

3. Höllerhage, M. (2025). Pesticides and parkinson’s disease: causal relationship at the population and individual level? Journal of Neural Transmission, 1–36.

4. Brown, T.P., Rumsby, P.C., Capleton, A.C., Rushton, L., and Levy, L.S. (2006). Pesticides and Parkinson’s disease--is there a link? Environ Health Perspect 114, 156–164. 10.1289/ehp.8095.

5. Pezzoli, G., and Cereda, E. (2013). Exposure to pesticides or solvents and risk of Parkinson disease. Neurology 80, 2035–2041.

6. Betarbet, R., Sherer, T.B., MacKenzie, G., Garcia-Osuna, M., Panov, A.V., and Greenamyre, J.T. (2000). Chronic systemic pesticide exposure reproduces features of Parkinson’s disease. Nature Neuroscience 3, 1301–1306. 10.1038/81834.

7. Greenamyre, J.T., Cannon, J.R., Drolet, R., and Mastroberardino, P.G. (2010). Lessons from the rotenone model of Parkinson’s disease. Trends Pharmacol Sci 31, 141–142; author reply 142-143. 10.1016/j.tips.2009.12.006.

8. Chia, S.J., Tan, E.-K., and Chao, Y.-X. (2020). Historical perspective: models of Parkinson’s disease. International journal of molecular sciences 21, 2464.

9. Innos, J., and Hickey, M.A. (2021). Using rotenone to model Parkinson’s disease in mice: a review of the role of pharmacokinetics. Chemical research in toxicology 34, 1223–1239.

10. Zuch, C.L., Nordstroem, V.K., Briedrick, L.A., Hoernig, G.R., Granholm, A.C., and Bickford, P.C. (2000). Time course of degenerative alterations in nigral dopaminergic neurons following a 6 - hydroxydopamine lesion. Journal of Comparative Neurology 427, 440–454.

11. Jackson-Lewis, V., Jakowec, M., Burke, R.E., and Przedborski, S. (1995). Time course and morphology of dopaminergic neuronal death caused by the neurotoxin 1-methyl-4-phenyl-1, 2, 3, 6-tetrahydropyridine. Neurodegeneration 4, 257–269.

12. Jeon, B.S., Jackson-Lewis, V., and Burke, R.E. (1995). 6-Hydroxydopamine lesion of the rat substantia nigra: time course and morphology of cell death. Neurodegeneration 4, 131–137.

13. Panov, A., Dikalov, S., Shalbuyeva, N., Taylor, G., Sherer, T., and Greenamyre, J.T. (2005). Rotenone model of Parkinson disease: multiple brain mitochondria dysfunctions after short term systemic rotenone intoxication. Journal of Biological Chemistry 280, 42026–42035.

14. Inden, M., Kitamura, Y., Abe, M., Tamaki, A., Takata, K., and Taniguchi, T. (2011). Parkinsonian rotenone mouse model: reevaluation of long-term administration of rotenone in C57BL/6 mice. Biological and Pharmaceutical Bulletin 34, 92–96.

15. Cannon, J.R., Tapias, V., Na, H.M., Honick, A.S., Drolet, R.E., and Greenamyre, J.T. (2009). A highly reproducible rotenone model of Parkinson’s disease. Neurobiology of Disease 34, 279–290. 10.1016/j.nbd.2009.01.016.

16. Van Laar, A.D., Webb, K.R., Keeney, M.T., Van Laar, V.S., Zharikov, A., Burton, E.A., Hastings, T.G., Glajch, K.E., Hirst, W.D., Greenamyre, J.T., and Rocha, E.M. (2023). Transient exposure to rotenone causes degeneration and progressive parkinsonian motor deficits, neuroinflammation, and synucleinopathy. NPJ Parkinsons Dis 9, 121. 10.1038/s41531-023-00561-6.

17. Sherer, T.B., Betarbet, R., Kim, J.-H., and Greenamyre, J.T. (2003). Selective microglial activation in the rat rotenone model of Parkinson’s disease. Neuroscience letters 341, 87–90.

18. Rocha, S.M., Bantle, C.M., Aboellail, T., Chatterjee, D., Smeyne, R.J., and Tjalkens, R.B. (2022). Rotenone induces regionally distinct α-synuclein protein aggregation and activation of glia prior to loss of dopaminergic neurons in C57Bl/6 mice. Neurobiology of disease 167, 105685.

19. Miyazaki, I., Isooka, N., Kikuoka, R., Imafuku, F., Masai, K., Tomimoto, K., Sakaguchi, M., Sogawa, C., Sogawa, N., and Kitamura, Y. (2025). Rotenone targets midbrain astrocytes to produce glial dysfunction-mediated dopaminergic neurodegeneration. Acta Neuropathologica Communications 13, 234.

20. Liu, Y., Sun, J.-D., Song, L.-K., Li, J., Chu, S.-F., Yuan, Y.-H., and Chen, N.-H. (2015). Environment-contact administration of rotenone: A new rodent model of Parkinson’s disease. Behavioural brain research 294, 149–161.

21. Morais, L.H., Hara, D.B., Bicca, M.A., Poli, A., and Takahashi, R.N. (2018). Early signs of colonic inflammation, intestinal dysfunction, and olfactory impairments in the rotenone-induced mouse model of Parkinson’s disease. Behavioural pharmacology 29, 199–210.

22. Mitra, S., Chakrabarti, N., Dutta, S., Ray, S., Bhattacharya, P., Sinha, P., and Bhattacharyya, A. (2015). Gender-specific brain regional variation of neurons, endogenous estrogen, neuroinflammation and glial cells during rotenone-induced mouse model of Parkinson’s disease. Neuroscience 292, 46–70.

23. De Miranda, B.R., Fazzari, M., Rocha, E.M., Castro, S., and Greenamyre, J.T. (2019). Sex Differences in Rotenone Sensitivity Reflect the Male-to-Female Ratio in Human Parkinson’s Disease Incidence. Toxicol Sci 170, 133–143. 10.1093/toxsci/kfz082.

24. Niederberger, E., Wilken-Schmitz, A., Manderscheid, C., Schreiber, Y., Gurke, R., and Tegeder, I. (2022). Non-reproducibility of oral rotenone as a model for Parkinson’s disease in mice. International journal of molecular sciences 23, 12658.

25. Poulin, J.F., Gaertner, Z., Moreno-Ramos, O.A., and Awatramani, R. (2020). Classification of Midbrain Dopamine Neurons Using Single-Cell Gene Expression Profiling Approaches. Trends Neurosci 43, 155–169. 10.1016/j.tins.2020.01.004.

26. Conrad, W.S., Oriol, L., Kollman, G.J., Faget, L., and Hnasko, T.S. (2024). Proportion and distribution of neurotransmitter-defined cell types in the ventral tegmental area and substantia nigra pars compacta. Addict Neurosci 13. 10.1016/j.addicn.2024.100183.

27. Poulin, J.F., Caronia, G., Hofer, C., Cui, Q., Helm, B., Ramakrishnan, C., Chan, C.S., Dombeck, D.A., Deisseroth, K., and Awatramani, R. (2018). Mapping projections of molecularly defined dopamine neuron subtypes using intersectional genetic approaches. Nat Neurosci 21, 1260–1271. 10.1038/s41593-018-0203-4.

28. Carmichael, K., Sullivan, B., Lopez, E., Sun, L., and Cai, H. (2021). Diverse midbrain dopaminergic neuron subtypes and implications for complex clinical symptoms of Parkinson’s disease. Ageing Neurodegener Dis 1. 10.20517/and.2021.07.

29. Pereira Luppi, M., Azcorra, M., Caronia-Brown, G., Poulin, J.-F., Gaertner, Z., Gatica, S., Moreno-Ramos, O.A., Nouri, N., Dubois, M., Ma, Y.C., et al. (2021). Sox6 expression distinguishes dorsally and ventrally biased dopamine neurons in the substantia nigra with distinctive properties and embryonic origins. Cell Reports 37, 109975. 10.1016/j.celrep.2021.109975.

30. Zhu, C., Vourc’h, P., Fernagut, P.O., Fleming, S.M., Lacan, S., Dicarlo, C.D., Seaman, R.L., and Chesselet, M.F. (2004). Variable effects of chronic subcutaneous administration of rotenone on striatal histology. Journal of Comparative Neurology 478, 418–426.

31. Tanguay, W., Ducrot, C., Giguère, N., Bourque, M.J., and Trudeau, L.E. (2021). Neonatal 6-OHDA lesion of the SNc induces striatal compensatory sprouting from surviving SNc dopaminergic neurons without VTA contribution. European Journal of Neuroscience 54, 6618–6632.

32. Lee, C.S., Sauer, H., and Björklund, A. (1996). Dopaminergic neuronal degeneration and motor impairments following axon terminal lesion by intrastriatal 6-hydroxydopamine in the rat. Neuroscience 72, 641–653. 10.1016/0306-4522(95)00571-4.

33. Stott, S.R., and Barker, R.A. (2014). Time course of dopamine neuron loss and glial response in the 6-OHDA striatal mouse model of P arkinson’s disease. European Journal of Neuroscience 39, 1042–1056.

34. Kozina, E., Khaindrava, V., Kudrin, V., Kucherianu, V., Klodt, P., Bocharov, E., Raevskiĭ, K., Kryzhanovskiĭ, G., and Ugriumov, M. (2010). Experimental modeling of functional deficiency of the nigrostriatal dopaminergic system in mice. Rossiiskii Fiziologicheskii Zhurnal Imeni IM Sechenova 96, 270–282.

35. Ugrumov, M., Khaindrava, V., Kozina, E., Kucheryanu, V., Bocharov, E., Kryzhanovsky, G., Kudrin, V., Narkevich, V., Klodt, P., and Rayevsky, K. (2011). Modeling of presymptomatic and symptomatic stages of parkinsonism in mice. Neuroscience 181, 175–188.

36. Li, L., Qin, H., Wang, J., Wang, J., Wang, X., and Gao, G. (2009). Axonal degeneration of nigra-striatum dopaminergic neurons induced by 1-methyl-4-phenyl-1, 2, 3, 6-tetrahydropyridine in mice. Journal of International Medical Research 37, 455–463.

37. Kenkhuis, B., Somarakis, A., Kleindouwel, L.R.T., van Roon-Mom, W.M.C., Höllt, T., and van der Weerd, L. (2022). Co-expression patterns of microglia markers Iba1, TMEM119 and P2RY12 in Alzheimer’s disease. Neurobiology of Disease 167, 105684. 10.1016/j.nbd.2022.105684.

38. Walker, D.G., Tang, T.M., Mendsaikhan, A., Tooyama, I., Serrano, G.E., Sue, L.I., Beach, T.G., and Lue, L.-F. (2020). Patterns of Expression of Purinergic Receptor P2RY12, a Putative Marker for Non-Activated Microglia, in Aged and Alzheimer’s Disease Brains. International Journal of Molecular Sciences 21, 678.

39. Iring, A., Tóth, A., Baranyi, M., Otrokocsi, L., Módis, L.V., Gölöncsér, F., Varga, B., Hortobágyi, T., Bereczki, D., Dénes, Á., and Sperlágh, B. (2022). The dualistic role of the purinergic P2Y12-receptor in an in vivo model of Parkinson’s disease: Signalling pathway and novel therapeutic targets. Pharmacological Research 176, 106045. 10.1016/j.phrs.2021.106045.

40. Sharma, N., and Nehru, B. (2015). Characterization of the lipopolysaccharide induced model of Parkinson’s disease: Role of oxidative stress and neuroinflammation. Neurochemistry International 87, 92–105.

41. Duffy, M.F., Collier, T.J., Patterson, J.R., Kemp, C.J., Luk, K.C., Tansey, M.G., Paumier, K.L., Kanaan, N.M., Fischer, D.L., Polinski, N.K., et al. (2018). Lewy body-like alpha-synuclein inclusions trigger reactive microgliosis prior to nigral degeneration. J Neuroinflammation 15, 129. 10.1186/s12974-018-1171-z.

42. Sanchez, H.L., Silva, L.B., Portiansky, E.L., Herenu, C.B., Goya, R.G., and Zuccolilli, G.O. (2008). Dopaminergic mesencephalic systems and behavioral performance in very old rats. Neuroscience 154, 1598–1606. 10.1016/j.neuroscience.2008.04.016.

43. Rudow, G., O’Brien, R., Savonenko, A.V., Resnick, S.M., Zonderman, A.B., Pletnikova, O., Marsh, L., Dawson, T.M., Crain, B.J., West, M.J., and Troncoso, J.C. (2008). Morphometry of the human substantia nigra in ageing and Parkinson’s disease. Acta Neuropathol 115, 461–470. 10.1007/s00401-008-0352-8.

44. Ma, S.Y., Röytt, M., Collan, Y., and Rinne, J.O. (1999). Unbiased morphometrical measurements show loss of pigmented nigral neurones with ageing. Neuropathol Appl Neurobiol 25, 394–399. 10.1046/j.1365-2990.1999.00202.x.

45. Wrangel, C.v., Schwabe, K., John, N., Krauss, J.K., and Alam, M. (2015). The rotenone-induced rat model of Parkinson’s disease: Behavioral and electrophysiological findings. Behavioural brain research 279, 52–61. 10.1016/j.bbr.2014.11.002.

46. Inden, M., Kitamura, Y., Takeuchi, H., Yanagida, T., Takata, K., Kobayashi, Y., Taniguchi, T., Yoshimoto, K., Kaneko, M., and Okuma, Y. (2007). Neurodegeneration of mouse nigrostriatal dopaminergic system induced by repeated oral administration of rotenone is prevented by 4-phenylbutyrate, a chemical chaperone. Journal of neurochemistry 101, 1491–1504.

47. Parameshwaran, K., Irwin, M.H., Steliou, K., and Pinkert, C.A. (2012). Protection by an antioxidant of rotenone-induced neuromotor decline, reactive oxygen species generation and cellular stress in mouse brain. Pharmacology Biochemistry and Behavior 101, 487–492.

48. Venkatesh Gobi, V., Rajasankar, S., Ramkumar, M., Dhanalakshmi, C., Manivasagam, T., Justin Thenmozhi, A., Essa, M.M., Chidambaram, R., and Kalandar, A. (2018). Agaricus blazei extract abrogates rotenone-induced dopamine depletion and motor deficits by its anti-oxidative and anti-inflammatory properties in Parkinsonic mice. Nutritional Neuroscience 21, 657–666.

49. Perez-Pardo, P., Broersen, L.M., Kliest, T., Van Wijk, N., Attali, A., Garssen, J., and Kraneveld, A.D. (2018). Additive effects of levodopa and a neurorestorative diet in a mouse model of Parkinson’s disease. Frontiers in Aging Neuroscience 10, 237.

50. Miyazaki, I., Isooka, N., Imafuku, F., Sun, J., Kikuoka, R., Furukawa, C., and Asanuma, M. (2020). Chronic systemic exposure to low-dose rotenone induced central and peripheral neuropathology and motor deficits in mice: reproducible animal model of Parkinson’s disease. International journal of molecular sciences 21, 3254.

51. Thomas Broome, S., and Castorina, A. (2022). Systemic rotenone administration causes extra-nigral alterations in C57BL/6 mice. Biomedicines 10, 3174.

52. Ming, G.-l., and Song, H. (2011). Adult neurogenesis in the mammalian brain: significant answers and significant questions. Neuron 70, 687–702.

53. Arzate, D.M., and Covarrubias, L. (2020). Adult neurogenesis in the context of brain repair and functional relevance. Stem cells and development 29, 544–554.

54. Gage, F.H. (2019). Adult neurogenesis in mammals. Science 364, 827–828.

55. Parent, J.M. (2003). Injury-induced neurogenesis in the adult mammalian brain. The Neuroscientist 9, 261–272.

56. Sun, D. (2014). The potential of endogenous neurogenesis for brain repair and regeneration following traumatic brain injury. Neural regeneration research 9, 688–692.

57. Varadarajan, S.G., Hunyara, J.L., Hamilton, N.R., Kolodkin, A.L., and Huberman, A.D. (2022). Central nervous system regeneration. Cell 185, 77–94.

58. Jin, Y., Dougherty, S.E., Wood, K., Sun, L., Cudmore, R.H., Abdalla, A., Kannan, G., Pletnikov, M., Hashemi, P., and Linden, D.J. (2016). Regrowth of serotonin axons in the adult mouse brain following injury. Neuron 91, 748–762.

59. Dougherty, S.E., Kajstura, T.J., Jin, Y., Chan-Cortés, M.H., Kota, A., and Linden, D.J. (2020). Catecholaminergic axons in the neocortex of adult mice regrow following brain injury. Experimental neurology 323, 113089.

60. Cooke, P., Janowitz, H., and Dougherty, S.E. (2022). Neuronal redevelopment and the regeneration of neuromodulatory axons in the adult mammalian central nervous system. Front. Cell. Neurosci. 16, 872501.

61. Burbulla, L.F., Song, P., Mazzulli, J.R., Zampese, E., Wong, Y.C., Jeon, S., Santos, D.P., Blanz, J., Obermaier, C.D., and Strojny, C. (2017). Dopamine oxidation mediates mitochondrial and lysosomal dysfunction in Parkinson’s disease. Science 357, 1255–1261.

62. Langston, R.G., Rudenko, I.N., Kumaran, R., Hauser, D.N., Kaganovich, A., Ponce, L.B., Mamais, A., Ndukwe, K., Dillman, A.A., Al-Saif, A.M., et al. (2019). Differences in Stability, Activity and Mutation Effects Between Human and Mouse Leucine-Rich Repeat Kinase 2. Neurochem Res 44, 1446–1459. 10.1007/s11064-018-2650-4.

63. Burns, T.C., Li, M.D., Mehta, S., Awad, A.J., and Morgan, A.A. (2015). Mouse models rarely mimic the transcriptome of human neurodegenerative diseases: A systematic bioinformatics-based critique of preclinical models. European journal of pharmacology 759, 101–117.

64. de Soysa, T.Y., Therrien, M., Walker, A.C., and Stevens, B. (2022). Redefining microglia states: Lessons and limits of human and mouse models to study microglia states in neurodegenerative diseases. (Elsevier), pp. 101651.

65. Li, J., Pan, L., Pembroke, W., Rexach, J., Godoy, M., Condro, M., Alvarado, A., Harteni, M., Chen, Y., and Stiles, L. (2021). Conservation and divergence of vulnerability and responses to stressors between human and mouse astrocytes. Nat Commun 12: 3958.

66. Fan, F., Funk, L., and Lou, X. (2016). Dynamin 1- and 3-Mediated Endocytosis Is Essential for the Development of a Large Central Synapse In Vivo. J Neurosci 36, 6097–6115. 10.1523/jneurosci.3804-15.2016.

67. Mahapatra, S., Fan, F., and Lou, X. (2016). Tissue-specific dynamin-1 deletion at the calyx of Held decreases short-term depression through a mechanism distinct from vesicle resupply. Proc Natl Acad Sci U S A 113, E3150–3158. 10.1073/pnas.1520937113.

68. Fan, F., Ji, C., Wu, Y., Ferguson, S.M., Tamarina, N., Philipson, L.H., and Lou, X. (2015). Dynamin 2 regulates biphasic insulin secretion and plasma glucose homeostasis. J Clin Invest 125, 4026–4041. 10.1172/jci80652.

69. Ji, C., Fan, F., and Lou, X. (2017). Vesicle Docking Is a Key Target of Local PI(4,5)P2 Metabolism in the Secretory Pathway of INS-1 Cells. Cell Rep 20, 1409–1421. 10.1016/j.celrep.2017.07.041.

70. Ji, C., and Lou, X. (2016). Single-molecule Super-resolution Imaging of Phosphatidylinositol 4,5-bisphosphate in the Plasma Membrane with Novel Fluorescent Probes. J Vis Exp. 10.3791/54466.

